# Genome-wide profiling reveals a dual role for histone H2A monoubiquitylation at Polycomb-repressed and enhancer chromatin

**DOI:** 10.1101/2023.05.28.542673

**Authors:** Kailynn MacGillivray, Daniel Fusca, Luomeng Tan, Reta Aram, Arneet L. Saltzman

## Abstract

Histone modifications are an integral component of eukaryotic genome regulation. Polycomb Repressive Complex 1 (PRC1) is responsible for depositing histone H2A lysine 119 monoubiquitylation (H2AK119ub) and can work cooperatively with PRC2-mediated histone H3 lysine 27 trimethylation (H3K27me3) to maintain gene repression. However, H3K27me3-independent functions and roles in gene activation have also been described for PRC1. Thus, the extent to which Polycomb complexes and their corresponding histone modifications function together or independently and the conservation of these roles in different organisms is unclear. Using *C. elegans* as a model, we investigated the relationship between H2AK119ub and H3K27me3. Here we show that the majority of H2AK119ub and H3K27me3 enrichment across the genome in embryos is distinct, and that the bulk levels of these modifications are regulated independently. We identify many genes related to nervous system development and functionality that have H2AK119ub-enriched promoters and are misregulated in H2AK119ub-deficient mutants, including a subset of genes that are normally H3K27me3-repressed. Surprisingly, we also find an enrichment of H2AK119ub at enhancers, including enhancers proximal to genes which are both up-regulated and down-regulated following the loss of this histone modification. Together, our results indicate a dual role for H2AK119ub in the regulation of both H3K27me3-repressed and enhancer chromatin states.

## Introduction

Polycomb Repressive Complexes are major regulators of chromatin state and gene expression. Polycomb proteins can be grouped into two families of multi-subunit complexes, <u>P</u>olycomb <u>R</u>epressive <u>C</u>omplex <u>2</u> (PRC2), which is responsible for methylation of histone H3K27 (H3K27me3), and <u>P</u>olycomb <u>R</u>epressive <u>C</u>omplex <u>1</u> (PRC1), which is responsible for monoubiquitylation of histone H2AK119 (H2AK119ub) (Kuroda et al. 2020). PRC1 complexes can be further sub-divided into canonical PRC1 (cPRC1) and variant PRC1 (vPRC1) based on their subunits and mechanism of genomic targeting (Blackledge et al. 2014; Chittock et al. 2017; Scelfo et al. 2019; Kuroda et al. 2020). In cPRC1, a Chromobox (CBX) protein subunit binds H3K27me3 via its chromodomain (Cao et al. 2002; Wang et al. 2004; Chittock et al. 2017; Scelfo et al. 2019; Kuroda et al. 2020). In vPRC1, RING1 and YY1 Binding Protein (RYBP) replaces the CBX subunit, and therefore targeting of this complex can be independent of H3K27me3 (Tavares et al. 2012; Morey et al. 2013; Blackledge et al. 2014; Chittock et al. 2017; Kuroda et al. 2020). Interestingly, vPRC1 complexes are more catalytically active than cPRC1 complexes *in vitro* and are responsible for the majority of H2AK119ub *in vivo*, potentially due to the ability of RYBP to stimulate ubiquitylation activity (Gao et al. 2012; Morey et al. 2013; Taherbhoy et al. 2015; Rose et al. 2016; Fursova et al. 2019; Kuroda et al. 2020). While the interaction between cPRC1 and H3K27me3 is well established, the relationship between vPRC1-deposited H2AK119ub and H3K27me3 is less well understood.

PRC1 activity has different impacts on gene expression depending on the context. PRC1 can induce H3K27me3-independent gene silencing via inhibition of Ser-5-phosphorylated RNA Polymerase II association with transcriptional start sites (Dobrinić et al. 2021). PRC1 can also function cooperatively and redundantly with PRC2 and H3K27me3 to mediate repression of target genes, such as the transcription factors *Klf4* and *Tbx3* (Cohen et al. 2018, 2021; Petracovici and Bonasio 2021; Sugishita et al. 2021). Variant PRC1 is involved in gene repression, but can also promote the expression of transcription factors involved in epidermal differentiation, *Satb1* and *Pou3f1* (Cohen et al. 2018). Additionally, cPRC1 components, including Psc/PCGF2,4, Pc/CBX, and Ph/PHC, occupy enhancer and promoter regions in *Drosophila* eye-antennal imaginal discs, and their binding correlates with increased expression of the associated genes and higher enhancer-promoter contact frequencies (Loubiere et al. 2020). Although the involvement of PRC1 in both gene repression and gene activation has been demonstrated, the role of H2AK119ub in each process is unresolved.

In *C. elegans*, the PRC1-like and PRC2 complexes are associated with mostly distinct developmental and fertility phenotypes. Mutations in *mig-32* or *spat-3,* the putative functional homologs of the core PRC1 components, PCGF and RING1A/B, respectively, greatly reduce levels of H2AK119ub. These mutant animals also have neuronal development defects, including defective migration, axon guidance, and posterior process termination of the hermaphrodite specific motor neurons (HSN neurons), PVQ neurons, and PLM neurons, respectively (Karakuzu et al. 2009; Pierce et al. 2018). In contrast, these defects have not been observed in mutants for the *C. elegans* PRC2 complex, composed of MES-2, MES-3, and MES-6. The *C. elegans* PRC2 complex is involved in *Hox* gene silencing and repression of transition genes in late embryos (Ross and Zarkower 2003; Yuzyuk et al. 2009). Its key role in maintaining germ cell identity and gene expression patterns leads to a characteristic maternal effect sterile (Mes) phenotype in PRC2 mutants (Gaydos et al. 2012; Patel et al. 2012). In contrast, there is currently no evidence for the involvement of the PRC1-like complex in these processes, as *mig-32* and *spat-3* mutant animals are fertile. Yet, both the PRC1-like and PRC2 complexes do act in the same genetic pathway to regulate male tail morphogenesis (Ross and Zarkower 2003; Karakuzu et al. 2009). Altogether, these phenotypes suggest that PRC2 and PRC1 play at least partially independent roles in *C. elegans* development. However, the target genes regulated by the *C. elegans* PRC1-like complex have not been identified, and it remains unclear to what extent these two complexes are functioning cooperatively or independently on a genome-wide level.

Here we use ChIP-seq and transcriptome profiling to investigate the gene regulatory roles of H2AK119ub in *C. elegans* embryos, capturing the developmental timepoint at which the PRC1 component homologs are most highly expressed (Hillier et al. 2009; Boeck et al. 2016). Our findings reveal that, in *C. elegans*, H2AK119ub peaks and H3K27me3 peaks are mostly distinct. Furthermore, while H2AK119ub is severely depleted in *mig-32* and *spat-3* mutants, H3K27me3 is not significantly affected. RNA-seq analysis in *C. elegans* embryos reveals that many genes with H2AK119ub-enriched promoters are differentially expressed in H2AK119ub-deficient mutants, a subset of which are involved in trans-synaptic signaling and are H3K27me3-repressed in wild-type animals. Surprisingly, we found that H2AK119ub is enriched at enhancer chromatin states along with enhancer-associated histone modifications, H3K4me1 and H3K27ac, including at enhancers proximal to genes that are differentially expressed following the loss of bulk H2AK119ub. Together, our results reveal the relationship between H2AK119ub and chromatin states and support the importance of PRC1-like complex activity in the regulation of *C. elegans* developmental gene expression.

## Results

### Genome-wide patterns of H2AK119ub and H3K27me3 are mostly distinct in *C. elegans* embryos

To investigate the relationship between H2AK119ub and H3K27me3 in *C. elegans,* we performed ChIP-seq for both histone modifications in embryos and examined their genomic distributions. A comparison of the consensus peaks, those called in two biological replicates, revealed that the majority of the H2AK119ub and H3K27me3 peaks do not overlap (Fig. 1A). Only 31% (1912/6098) of the H2AK119ub consensus peaks overlap by at least 50% of their length with H3K27me3 consensus peaks (Fig. 1B). Likewise, 17% (969/5687) of the H3K27me3 consensus peaks overlap by at least 50% of their length with H2AK119ub consensus peaks (Fig. 1B). To further compare the genomic distributions of these two histone modifications, we examined their distributions over whole chromosomes (Fig. 1C). In *C. elegans*, H3K27me3 is enriched at the heterochromatin-rich chromosome arms (Liu et al. 2011), particularly the arms of chromosome II and chromosome V (Fig. 1C). However, unexpectedly and in contrast to H3K27me3, H2AK119ub is not enriched on the chromosome arms. These peak and distribution patterns suggest that PRC1-mediated H2AK119ub has functions that are separate from H3K27me3-marked heterochromatin and Polycomb domains. As well, the distinct distributions of these two histone modifications are in contrast to the co-enrichment of PRC1 and PRC2 previously observed in mouse embryonic stem cells, mammalian epidermal progenitor cells and *Drosophila* (Boyer et al. 2006; Schwartz et al. 2006; Cohen et al. 2018).

**Figure 1.**
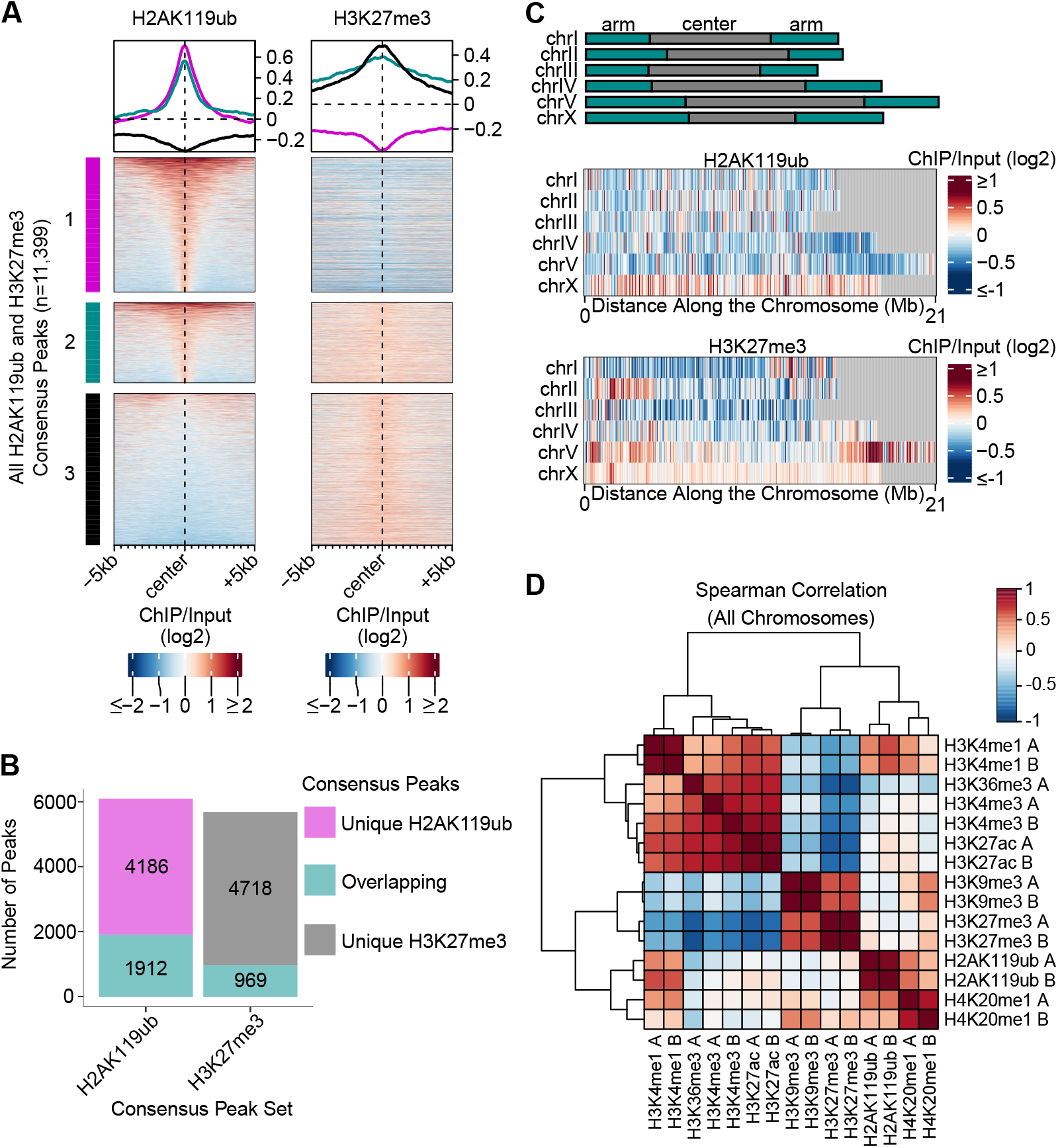
H2AK119ub and H3K27me3 have distinct genomic distributions in *C. elegans* embryos. **A.** Heatmaps of H2AK119ub and H3K27me3 ChIP-seq (log2-transformed ChIP/Input ratios) at all H2AK119ub and H3K27me3 consensus peaks. Consensus peaks are those called in two biological replicates. Peaks are organized into clusters, with cluster 1 containing the distinct H2AK119ub consensus peaks. Cluster 2 contains H2AK119ub and H3K27me3 peaks that overlap by at least 50% of their lengths. Cluster 3 contains the distinct H3K27me3 consensus peaks. **B.** Quantification of the overlap of H2AK119ub and H3K27me3 consensus peaks shown in A. Peaks are called as overlapping if at least 50% of their peak region overlaps with the other. **C.** Heatmaps of H2AK119ub and H3K27me3 ChIP-seq (log2-transformed ChIP/Input ratios) across whole *C. elegans* chromosomes. Scale represents average enrichment in 50 kb bins. The representative diagram indicates the positions of the arms and centers of each chromosome (top) (Rockman and Kruglyak 2009). **D.** Heatmap of the Spearman Correlation coefficients for the distributions of different histone modifications across all *C. elegans* chromosomes. Rows and columns are organized by complete Euclidean hierarchical clustering. A and B indicate biological replicates.

To compare the enrichment patterns of H2AK119ub to other histone modifications, we used ChIP-seq data from the modEncode consortium (Ho et al. 2014), computed pairwise Spearman Correlation coefficients for coverage across the genome (Fig. 1D) and compared enrichment across chromosomes (Supplemental Fig. S1). On a genome-wide scale, the heterochromatin-associated marks H3K27me3 and H3K9me3 are highly correlated, as previously described (Ho et al. 2014), while H2AK119ub has a very weak overall correlation with H3K27me3 (Fig. 1D). Instead, H2AK119ub is positively correlated with H3K4me1 (Fig. 1D), a mark associated with enhancers and promoters (Heintzman et al. 2009; Ho et al. 2014), and with H4K20me1, a mark enriched on the X chromosome (Supplemental Fig. S1) by the dosage compensation complex (Vielle et al. 2012; Wells et al. 2012; Kramer et al. 2015; Brejc et al. 2017). Notably, although the PRC1-like complex has not been previously implicated in dosage compensation, H2AK119ub is unexpectedly enriched on the X-chromosome (Fig. 1C, Supplemental Fig. S1), which likely accounts for its correlation with H4K20me1. Our results therefore suggest that H2AK119ub may play a role in X-chromosome regulation, potentially in conjunction with either PRC2-deposited H3K27me3 (Fong et al. 2002; Bender et al. 2004) or with the dosage compensation complex. However, across the genome, the distribution of H2AK119ub has a low correlation with H3K27me3, which suggests that these two histone modifications play independent roles in gene regulation and are associated with distinct chromatin states.

Given the unexpected distribution of H2AK119ub, we investigated whether recognition of the monoubiquitylated form of the H2A variant H2A.Z (*C. elegans* HTZ-1) in our ChIP-seq assays could account for this pattern (Supplemental Fig. S2). When comparing our H2AK119ub ChIP-seq with previously published HTZ-1 ChIP-seq (Whittle et al. 2008), only 9% (551/6098) of H2AK119ub consensus peaks overlap with HTZ-1 peaks (Supplemental Fig. S2B, C). Furthermore, previous profiling revealed an enrichment of HTZ-1 across gene bodies (Latorre et al. 2015), which is distinct from the distribution of H2AK119ub at upstream regions in our data (Fig. 2A). Our analysis therefore supports that the vast majority of the H2AK119ub ChIP-seq signal is from H2AK119ub and not HTZ-1ub.

**Figure 2.**
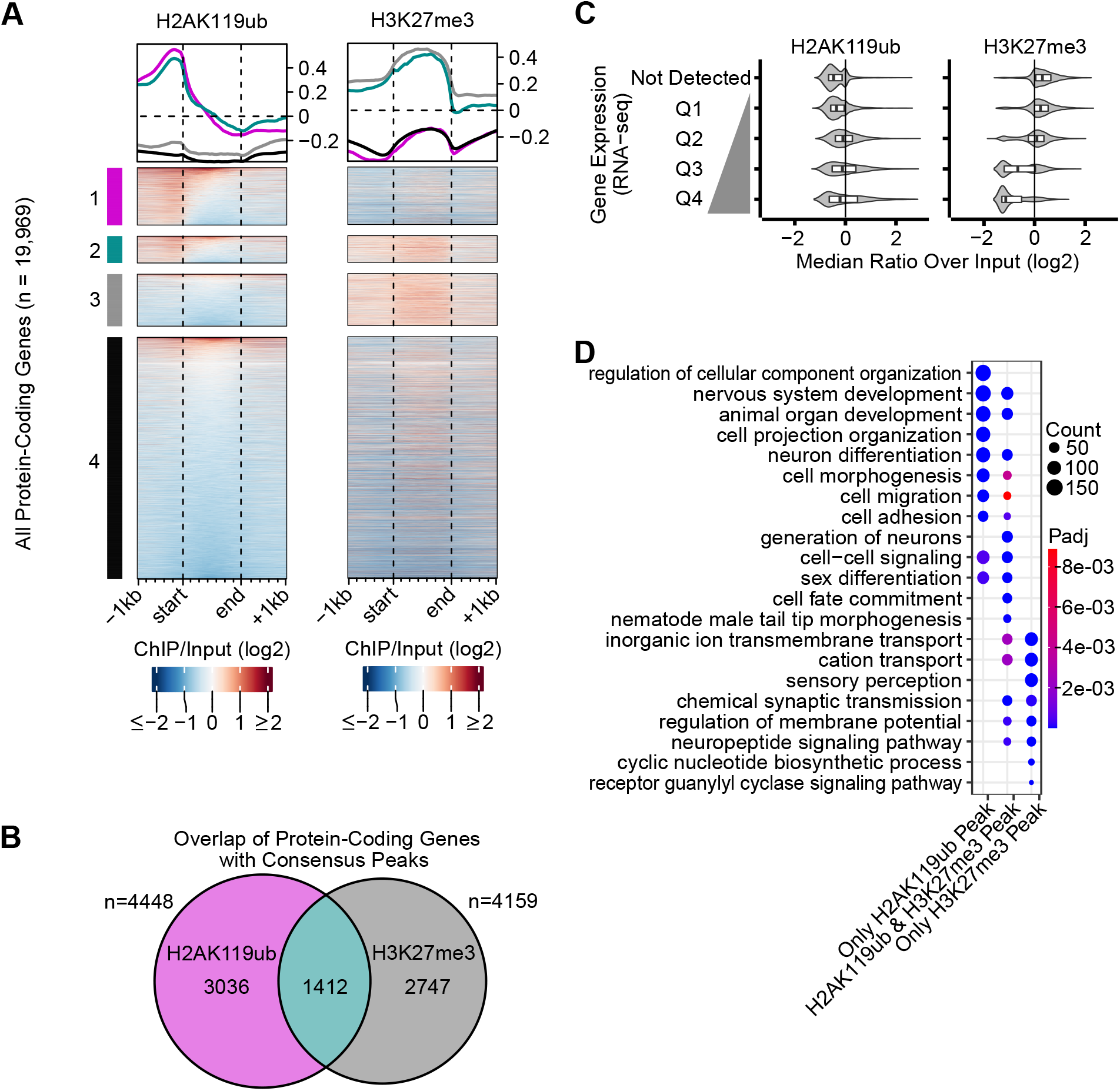
H2AK119ub and H3K27me3 are enriched at different protein-coding genes. **A.** Heatmaps of H2AK119ub and H3K27me3 ChIP-seq (log2-transformed ChIP/Input ratios) at all *C. elegans* protein-coding genes and flanking regions. The protein-coding genes are divided into 4 clusters, depending on their annotation with associated peaks: cluster 1: H2AK119ub only, cluster 2: both H2AK119ub and H3K27me3, cluster 3: H3K27me3 only, and cluster 4: neither an H2AK119ub nor an H3K27me3 peak. **B.** Quantification of the overlap between genes associated with H2AK119ub and/or H3K27me3 from clusters 1-3 in panel A. **C.** Violin plots of the median H2AK119ub or H3K27me3 signal (log2-transformed ChIP/Input ratio) at the promoters of protein-coding genes divided into quartiles of expression based on RNA-seq data in N2 (wild-type) embryos. H3K27me3, but not H2AK119ub is enriched over promoters of silent or lowly expressed genes. **D.** Enriched GO terms associated with genes with an H2AK119ub but no H3K27me3 peak, genes with both an H2AK119ub and H3K27me3 peak, or genes with an H3K27me3 but no H2AK119ub peak. The gene sets correspond to the genes in clusters 1-3 of panel A.

To determine if the pattern of distinct H2AK119ub and H3K27me3 peaks is conserved in mammals, we used previously generated ChIP-seq data to look at the enrichment of these two histone modifications at all H2AK119ub peaks in human induced pluripotent stem cells (iPSCs) (Chan et al. 2018), mouse epidermal progenitor cells (Cohen et al. 2018), leukemia cell line K562 (van den Boom et al. 2016), and breast cancer cell line T47D (Chan et al. 2018) (Supplemental Fig. S3). In all cell types examined, we observed a subset of H2AK119ub peaks that are co-enriched with H3K27me3, whereas others are distinct from H3K27me3 enrichment (Supplemental Fig. S3). These findings are consistent with the known co-targeting of repressed genes by PRC1 and PRC2 complexes in mammals (Boyer et al. 2006; Cohen et al. 2018), but also indicate that, in some genomic contexts, H2AK119ub functions independently from H3K27me3. Furthermore, the pattern of distinct H2AK119ub peaks is not unique to *C. elegans*, as regions enriched for H2AK119ub but not H3K27me3 are also observed in mammalian cells.

### H2AK119ub and H3K27me3 are associated with different sets of protein-coding genes

To determine if the distinct genomic distributions of H2AK119ub and H3K27me3 peaks in *C. elegans* extends to potential target genes, we examined the distribution of these modifications over all protein-coding genes and their flanking regions (Fig. 2A). Generally, H2AK119ub is most enriched over upstream promoter regions, whereas H3K27me3 enrichment extends from upstream to downstream regions, with the highest enrichment over the gene body (Fig. 2A). Of the protein-coding genes with associated H2AK119ub or H3K27me3 consensus peaks (n=7195), only 20% (n=1412) are annotated with both (Fig. 2B), which nevertheless represents a higher overlap than would be expected by chance (p < 2.2e-16, Fisher’s exact test). The enrichment of H2AK119ub and H3K27me3 at genes is consistent with the genome-wide patterns above, and supports regulation of mostly distinct sets of genes, with a smaller subset of common targets.

To compare these gene sets, we examined their chromosomal locations and category of expression pattern (Supplemental Fig. S4) based on previously defined groups (Knutson et al. 2016). Genes with H2AK119ub peaks, H3K27me3 peaks, or both H2AK119ub and H3K27me3 peaks are enriched in the ‘soma-specific’ gene expression category and on the X-chromosome and are depleted from the ‘germline-enriched’ category (Supplemental Fig. S4). In contrast, genes uniquely associated with H2AK119ub but not H3K27me3 peaks are enriched in the ‘ubiquitous’ expression category (Supplemental Fig. S4C). These patterns are consistent with the enrichment of H2AK119ub on the X-chromosome (Fig. 1C, Supplemental Fig. S1) and the previously reported depletion of germline-expressed genes from the X-chromosome (Gaydos et al. 2012), and also suggest that H2AK119ub regulates broadly expressed genes.

To further compare the expression levels of genes associated with H2AK119ub and H3K27me3, we looked at the enrichment of these two histone modifications over the promoters of protein-coding genes divided into different quartiles of expression based on RNA-seq data in N2 (wild-type) embryos (Fig. 2C). Consistent with its known role in gene repression, H3K27me3 is enriched over the promoters of silent and lowly expressed genes and is depleted from the promoters of genes in the highest quartiles of expression (Fig. 2C). In contrast, H2AK119ub does not show this pattern and is not enriched at lowly expressed or silent genes (Fig. 2C). Thus, H2AK119ub shows less association with gene repression than H3K27me3, suggesting that the two histone modifications play distinct roles in the regulation of gene expression.

To determine if the H2AK119ub-enriched and H3K27me3-enriched protein-coding genes are associated with different biological functions, we performed gene ontology (GO) enrichment analysis. The genes with an H2AK119ub peak are associated with nervous system development, neuron differentiation, and cell projection morphogenesis (Fig. 2D). This enrichment is consistent with roles for the putative *C. elegans* PRC1 subunit homologues *mig-32* and *spat-3* in neuronal migration, axon guidance, and neuronal cell fate (Karakuzu et al. 2009; Pierce et al. 2018; Bordet et al. 2022). On the other hand, H3K27me3-enriched genes are involved in neuropeptide signaling, sensory perception, and cyclic nucleotide biosynthesis (Fig. 2D). Genes with both H2AK119ub and H3K27me3 peaks share GO terms with genes associated with only one of the two modifications. However, these co-enriched genes are also uniquely associated with male tail tip morphogenesis, which is consistent with defects in sensory ray development of the male tail observed in mutants for both *C. elegans* PRC2 and putative PRC1 subunit homologs (Ross and Zarkower 2003; Karakuzu et al. 2009). In sum, H2AK119ub and H3K27me3 are associated with mostly distinct sets of protein-coding genes which are involved in different biological processes.

### H2AK119ub and H3K27me3 are not dependent on each other’s presence

To investigate if H2AK119ub and H3K27me3 are interdependent, we used western blotting to look at the bulk levels of these histone marks in mutants of putative PRC1 component homologs, *mig-32* and *spat-3*. The *mig-32(n4275)* allele is a deletion spanning the start of the coding region and the *spat-3(mgw26)* allele is a deletion of the entire coding region (Supplemental Fig. S5, Table S1). The *spat-3(gk22)* allele is a deletion that does not overlap with the conserved RING domain (Supplemental Fig. S5B) and is likely a partial loss-of-function. In *spat-3(gk22)* mutants, H2AK119ub is reduced to ∼60% of wild-type levels, and in *mig-32(n4275), spat-3(mgw26), mig-32(n4275)spat-3(gk22),* and *mig-32(n4275);spat-3(mgw26)* mutants, H2AK119ub levels are drastically reduced to 2-5% of wild-type (Fig. 3A). Strikingly, although H2AK119ub is severely depleted in the *mig-32(n4275)* and *spat-3(mgw26)* mutants, H3K27me3 levels are not reduced (Fig. 3A).

**Figure 3.**
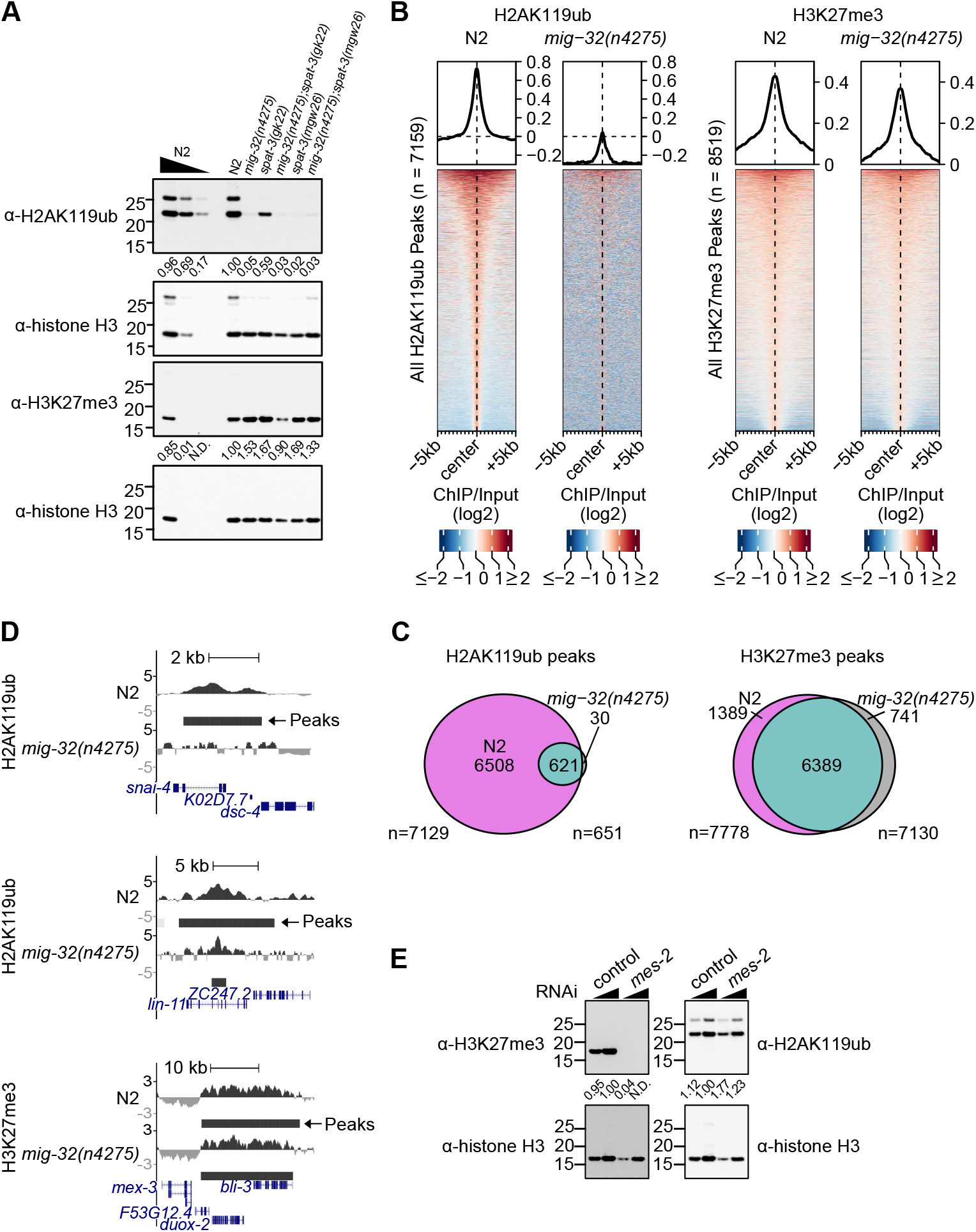
Levels of H2AK119ub and H3K27me3 are independent in *C. elegans* embryos. **A.** Western blot for H2AK119ub and H3K27me3 in *mig-32* and *spat-3* single and double mutants. Histone H3 was probed as a control. H2AK119ub, but not H3K27me3, is depleted in the mutants. The predicted relative migration distances are 23 kDa for H2AK119ub and 17 kDa for H3K27me3 and histone H3. See Supplemental Figure S5 and Table S1 for allele details. N.D., not detectable. **B.** Heatmaps of H2AK119ub and H3K27me3 (log2-transformed ChIP/Input ratios) at all H2AK119ub and H3K27me3 peaks in N2 (wild-type) and *mig-32(n4275)* deletion mutants. **C.** Quantification of the number of H2AK119ub (left) and H3K27me3 (right) peaks in N2 versus *mig-32(n4275)* mutants. The majority of H2AK119ub, but not H3K27me3, is lost in the mutants. **D.** UCSC Genome Browser view of a representative H2AK119ub peak that is lost in the *mig-32(n4275)* mutants (top), an H2AK119ub peak that is attenuated in the *mig-32(n4275)* mutants (middle), and an H3K27me3 peak that is retained in both genotypes (bottom). Coverage is displayed as the log2-transformed ChIP/Input ratio. Positions of MACS2-predicted peaks are shown below with darker colours indicating higher confidence. **E.** Western blot for H2AK119ub and H3K27me3 in embryos collected from adults exposed to control or *mes-2* RNAi. Histone H3 was probed as a control. H3K27me3, but not H2AK119ub, is depleted in the *mes-2* knockdown embryos. N.D., not detectable.

To determine if this pattern is also observed at the level of peaks, we performed ChIP-seq in wild-type (N2) and *mig-32(n4275)* embryos. As expected, the majority (91%; 6508/7129) of H2AK119ub peaks called in N2 were lost in *mig-32(n4275)* mutants (Fig. 3B,C). For example, the H2AK119ub peak over the gene body of *snai-1* and *dsc-4* is lost in *mig-32(n4275)* mutants (Fig. 3D, top panel). The reduction of H2AK119ub in *mig-32(n4275)* mutants was further validated by ChIP-qPCR at three representative peaks (Supplemental Fig. S6A,B). Moreover, the remaining 8.7% (621/7129) of H2AK119ub peaks called as common between the N2 and *mig-32(n4275)* genotypes were generally attenuated and substantially shorter in the mutants (Fig. 3B, Supplemental Fig. S6C), such as the peak in the gene body of *lin-11* and upstream of *ZC247.2* (Fig. 3D, middle panel). The genes associated with the attenuated H2AK119ub peaks called in *mig-32(n4275)* mutants did not appear to be a functionally distinct subset of genes, since they had similar GO terms to the genes associated with H2AK119ub peaks in wild-type animals (Supplemental Fig. S6D, Fig. 2D). In contrast to the loss of H2AK119ub, heatmaps of H3K27me3 peaks in N2 and *mig-32(n4275)* mutants are nearly indistinguishable, and the vast majority (82%; 6389/7778) of the H3K27me3 peaks are common to both N2 and the *mig-32(n4275)* mutants (Fig. 3B,C,D, bottom panel). Furthermore, the N2-specific H3K27me3 peaks are not enriched for H2AK119ub (Supplemental Fig. S6E). These results suggest that the deposition and genomic distribution of H3K27me3 is not dependent on *mig-32* or H2AK119ub.

To test if the converse is also true, if H2AK119ub deposition depends on PRC2/H3K27me3, we used western blotting to assess the levels of H2AK119ub and H3K27me3 following knockdown of *mes-2*, the *C. elegans* homolog of *E(z)*, the H3K27 methyltransferase of the *Drosophila* PRC2 complex (Bender et al. 2004). As expected, in embryos collected from adults exposed to *mes-2* RNAi, H3K27me3 is severely depleted (Fig. 3E). However, the levels of H2AK119ub are not disrupted (Fig. 3E), indicating that the bulk level of H2AK119ub is not dependent on the presence of H3K27me3. Together, these results indicate that the two histone modifications are predominantly regulated independently.

### Loss of H2AK119ub results in a larger magnitude of up-regulation than down-regulation of gene expression

To investigate the role of H2AK119ub in the regulation of gene expression, we performed RNA-seq in N2 (wild-type), *mig-32(n4275)*, *spat-3(mgw26)*, and *mig-32(n4275);spat-3(mgw26)* late embryos (Fig. 4, Supplemental Fig. S7). Late embryos were used to capture the timeframe in which *mig-32* and *spat-3* are most highly expressed in N2 animals (Hillier et al. 2009; Boeck et al. 2016). To classify genes as differentially expressed, we used a statistical significance threshold (adjusted p-value ≤ 0.05) and evaluated two different fold-change thresholds. At a 2-fold-change cut-off (absolute log2 fold-change of 1), 81-94% of the differentially expressed genes are up-regulated and 6-13% are down-regulated, depending on the genotype (Fig. 4A). At a 1.5-fold-change cut-off, 40-70% of the differentially expressed genes are up-regulated and 30-60% are down-regulated (Fig. 4A). Therefore, the loss of H2AK119ub has a larger impact on gene up-regulation in terms of magnitude. In the analyses below, we use a 1.5-fold change cut-off so as not to exclude the cohort of genes (892/1283) with modest but statistically significant changes, in particular because such changes may reflect differences in specific cell types that are partially masked when examining the embryos as a whole. Moreover, this cut-off has been used previously to identify genes differentially expressed following manipulation of Polycomb complexes (Jaensch et al. 2021; Petracovici and Bonasio 2021).

**Figure 4.**
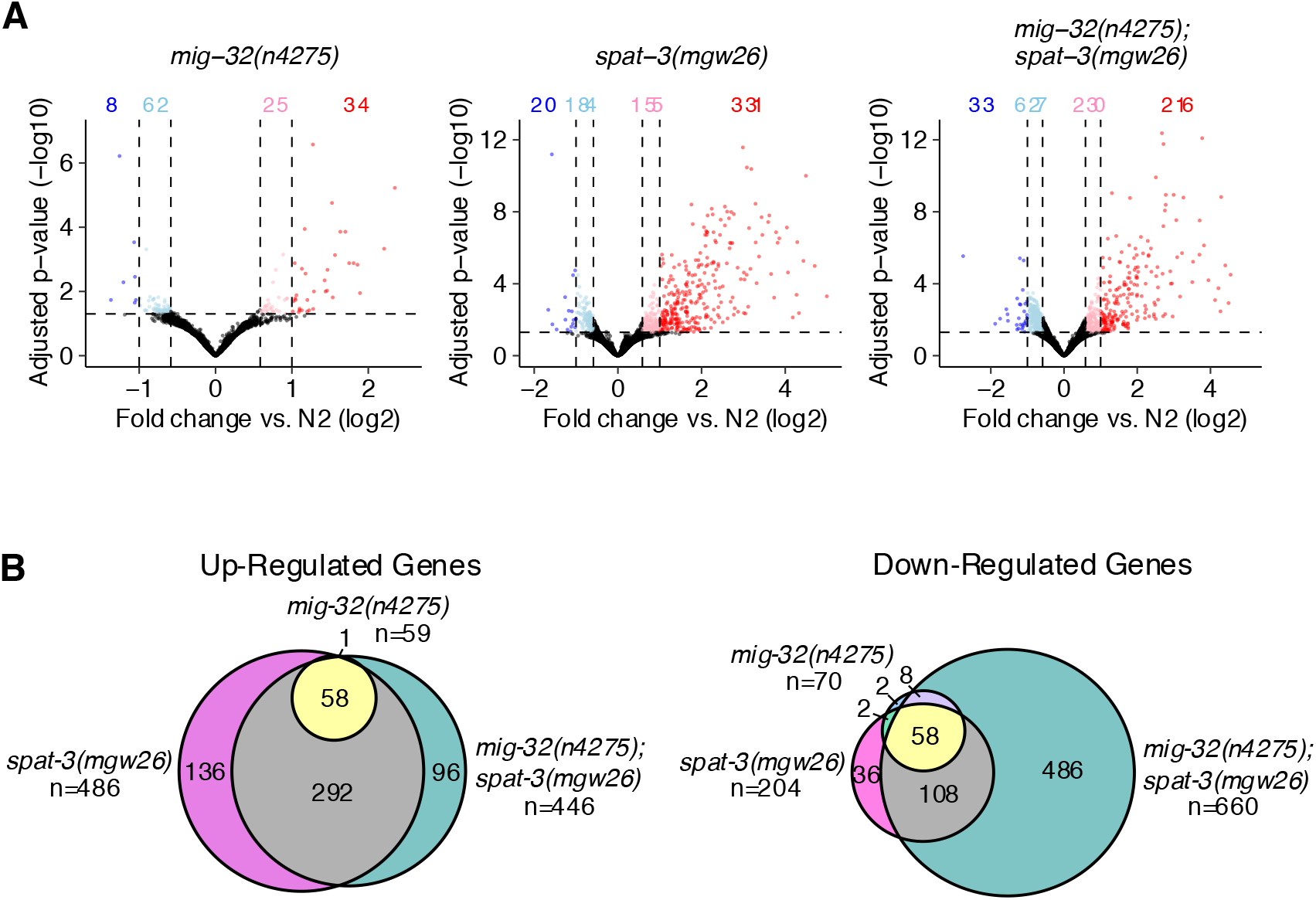
Differential gene expression in H2AK119ub-deficient mutants. **A.** Volcano plots of RNA-seq data in *mig-32(n4275)*, *spat-3(mgw26)*, and *mig-32(n4275);spat-3(mgw26)* mutants. Significantly differentially expressed genes (p-adjust ≤ 0.05) with an absolute fold change of at least 2 (log2 fold change of 1) are marked in red (upregulated) or blue (downregulated). Significantly differentially expressed genes with an absolute fold change of at least 1.5 (absolute log2 fold change of 0.58) are marked in pink (upregulated) or light blue (downregulated). **B.** Quantification of the overlap of the significantly up-regulated genes (left) and the significantly down-regulated genes (right) between the different mutant genotypes.

The *mig-32(n4275)* expression profile was more similar to N2 than the other mutants, with fewer differentially expressed genes (Fig. 4A, Supplemental Fig. S7B,C). The *spat-3(mgw26)* and *mig-32(n4275);spat-3(mgw26)* mutants share 78% (352/449) of their up-regulated genes (Fig. 4B). Of these commonly up-regulated genes, 13% (58/449) are also up-regulated in the *mig-32(n4275)* mutants, suggesting that the other 65% (294/449) of protein-coding genes are dependent on SPAT-3, but not MIG-32, for appropriate expression (Fig. 4B). Most of the down-regulated genes in the *mig-32(n4275)* mutants are also down-regulated in the *spat-3(mgw26)* single mutants (Fig. 4B). However, there are substantially more genes with a decrease in expression in the *mig-32(n4275);spat-3(mgw26)* genotype (Fig. 4A). Amongst the set of significantly, but modestly, down-regulated genes in the double mutants, 73% (492/670) are uniquely differentially expressed in the *mig-32(n4275);spat-3(mgw26)* double mutants, suggesting that MIG-32 and SPAT-3 may be working redundantly to promote the expression of these genes (Fig. 4B).

### Genes mis-regulated in H2AK119ub-deficient mutants are enriched for nervous system functions

To determine how the genes that are differentially expressed in any of the mutant genotypes correlate with chromatin states and what biological processes they may be involved in, we plotted the enrichment of H2AK119ub and H3K27me3 across their gene bodies and flanking regions and tested for enriched GO terms (Fig. 5A, B). Overall, 41% (524/1283) of the differentially expressed genes are associated with H2AK119ub and/or H3K27me3 peaks (Fig. 5C). We grouped the differentially expressed genes into 3 clusters based on the pattern of proximal H2AK119ub and H3K27me3 in N2 (wild-type), as well as their change in transcript level in H2AK119ub-deficient mutants versus N2 (Fig. 5A). Cluster 1 contains genes that have high H2AK119ub enrichment, particularly over their promoter regions (Fig. 5A). Cluster 2 contains genes with weak H2AK119ub enrichment over their promoters, but high H3K27me3 across their gene bodies. The cluster 2 genes are lowly expressed in N2 animals but 81% (474/587) are up-regulated in the H2AK119ub-deficient mutants (Fig. 5A). The genes in cluster 1 are associated with GO terms involved in nervous system development and neuron differentiation, which is consistent with GO terms associated with all protein-coding genes with H2AK119ub peaks (Fig. 5B, Fig. 2D). Interestingly, two-thirds of the genes in cluster 1 associated with these GO terms are down-regulated in the H2AK119ub-deficient mutants (Fig. 5A, B), suggesting that H2AK119ub, SPAT-3, or both may be promoting their expression in N2 animals. Similarly, the genes in cluster 2 are enriched for GO terms involved in trans-synaptic signaling, neurotransmitter transport and secretion (Fig. 5B). The differentially expressed genes in cluster 1 and cluster 2 are likely to contribute to the defects in neuron migration and axon guidance observed in the *mig-32* and *spat-3* mutants (Karakuzu et al. 2009; Pierce et al. 2018).

**Figure 5.**
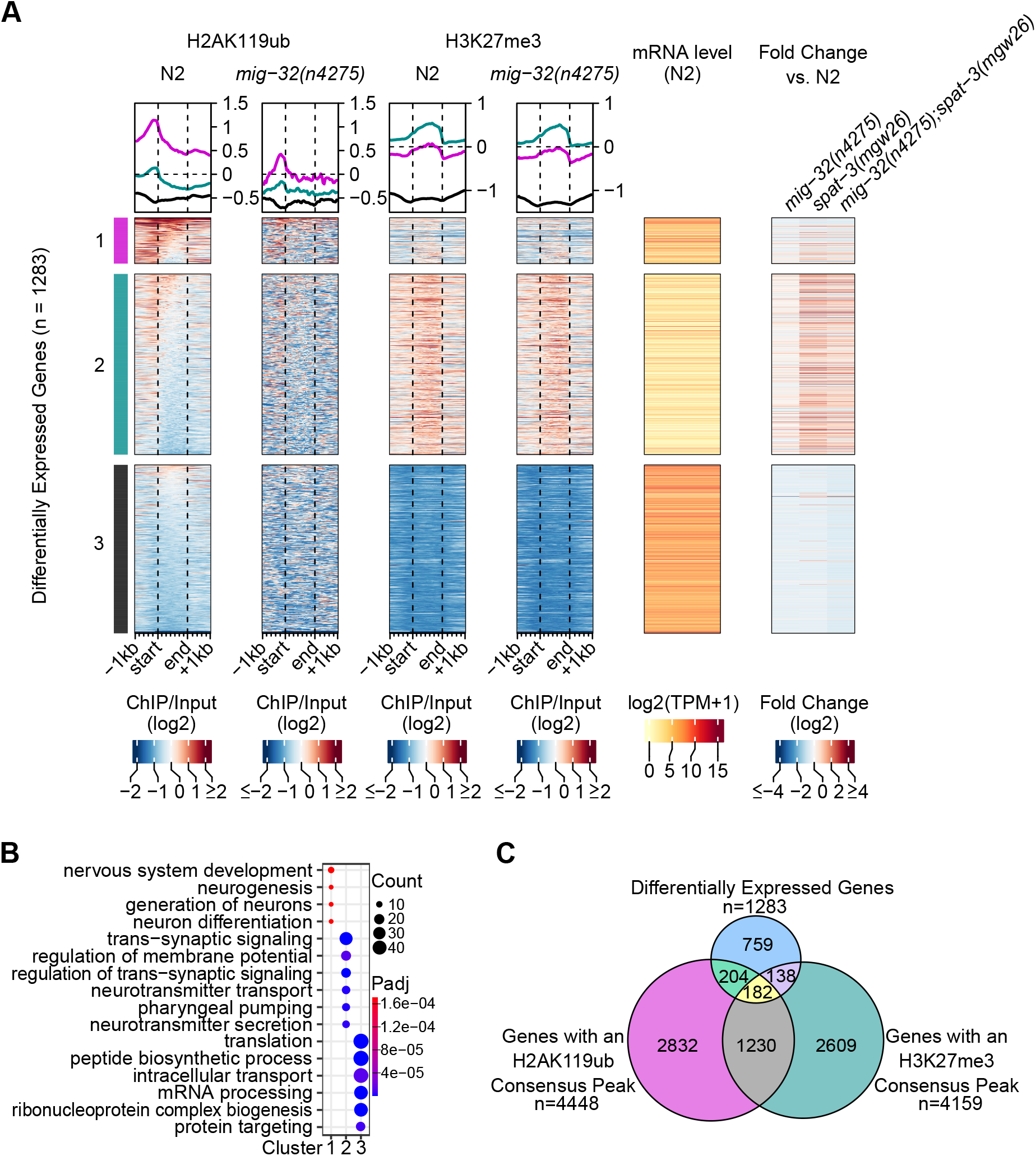
A subset of genes with H2AK119ub-enriched promoters are differentially expressed in *mig-32(4275)* and *spat-3(mgw26)* mutants. **A.** Heatmaps of H2AK119ub and H3K27me3 (log2-transformed ChIP/Input ratios) at the differentially expressed protein-coding genes in N2 (wild-type) and *mig-32(n4275)* mutants. N2 mRNA level in transcripts per million (TPM) and the fold change (log2) in each mutant genotype is displayed. The differentially expressed genes are grouped into 3 k-means clusters based on N2 H2AK119ub and H3K27me3 enrichment as well as fold change. **B.** Enriched gene ontology (GO) biological process terms associated with the differentially expressed protein-coding genes in each of the k-means clusters in panel A. **C.** Quantification of the overlap of protein-coding genes annotated with H2AK119ub consensus peaks, H3K27me3 consensus peaks, or called as significantly differentially expressed in H2AK119ub-deficient mutants.

Cluster 3 contains genes which do not show enrichment for H2AK119ub or H3K27me3 (Fig. 5A). These genes are highly expressed in N2 embryos but are slightly down-regulated in the H2AK119ub-deficient mutants, and are related to translation, mRNA processing, and protein targeting (Fig. 5A, B). Since these genes are down-regulated in the *mig-32* and *spat-3* mutants, particularly in the double mutants, the *C. elegans* PRC1-like complex may promote their expression. However, considering that there is no proximal H2AK119ub enrichment, promotion of the expression of these genes may be an indirect effect or may be mediated by H2AK119ub-independent activities.

Although many H2AK119ub-enriched protein-coding genes are differentially expressed following the loss of this histone modification, not all H2AK119ub-enriched genes are impacted (Fig. 5C, Supplemental Fig. S8). At a 1.5-fold-change cut-off, only 8.7% (386/4448) of the protein-coding genes annotated with an H2AK119ub peak are differentially expressed in the *mig-32(n4275)* and *spat-3(mgw26)* mutants (Fig. 5C). Likewise, at a 2-fold-change cut-off, only 3.8% (167/4448) of the H2AK119ub-enriched genes are differentially expressed (Supplemental Fig. S8B). These results suggest that many PRC1-like targets may be regulated by an H2AK119ub-independent mechanism or that the role of H2AK119ub is redundant in regulating the expression of these genes. Furthermore, only 13% (182/1412) of the protein-coding genes with both H2AK119ub and H3K27me3 peaks have an expression level change of at least 1.5 fold and only 6.4% have a fold change of at least 2 in the H2AK119ub-deficient mutants (Fig. 5C, Supplemental Fig. S8B), implying that the non-differentially expressed co-enriched genes may be dependent only on H3K27me3 for proper expression or may be regulated redundantly by the PRC1-like and PRC2 complexes. Nonetheless, the different chromatin states associated with each cluster of differentially expressed genes supports both H3K27me3-dependent and H3K27me3-independent activities of a PRC1-like complex in *C. elegans*.

### H2AK119ub is enriched at enhancer chromatin states

To look for a relationship between H2AK119ub and other histone modifications, we plotted the H2AK119ub ChIP-seq signal over different chromatin states (Fig. 6A). We used two sets of Hidden Markov Model (HMM)-predicted *C. elegans* chromatin states, one with 7 predicted chromatin states based on 8 histone modification ChIP-seq datasets (Daugherty et al. 2017), and one with 20 predicted chromatin states based on 17 histone and histone modification ChIP-chip or ChIP-seq datasets (Evans et al. 2016). Surprisingly, the H2AK119ub ChIP-seq signal is consistently highest at enhancer chromatin states, including predicted repressed and active enhancers, intergenic, intronic, and weak enhancers (Fig. 6A). In addition, histone modifications associated with enhancers, H3K4me1 and H3K27ac (Heintzman et al. 2009; Creyghton et al. 2010; Rada-Iglesias et al. 2011; Ho et al. 2014), are strongly enriched at the H2AK119ub consensus peaks (Fig. 6B).

**Figure 6.**
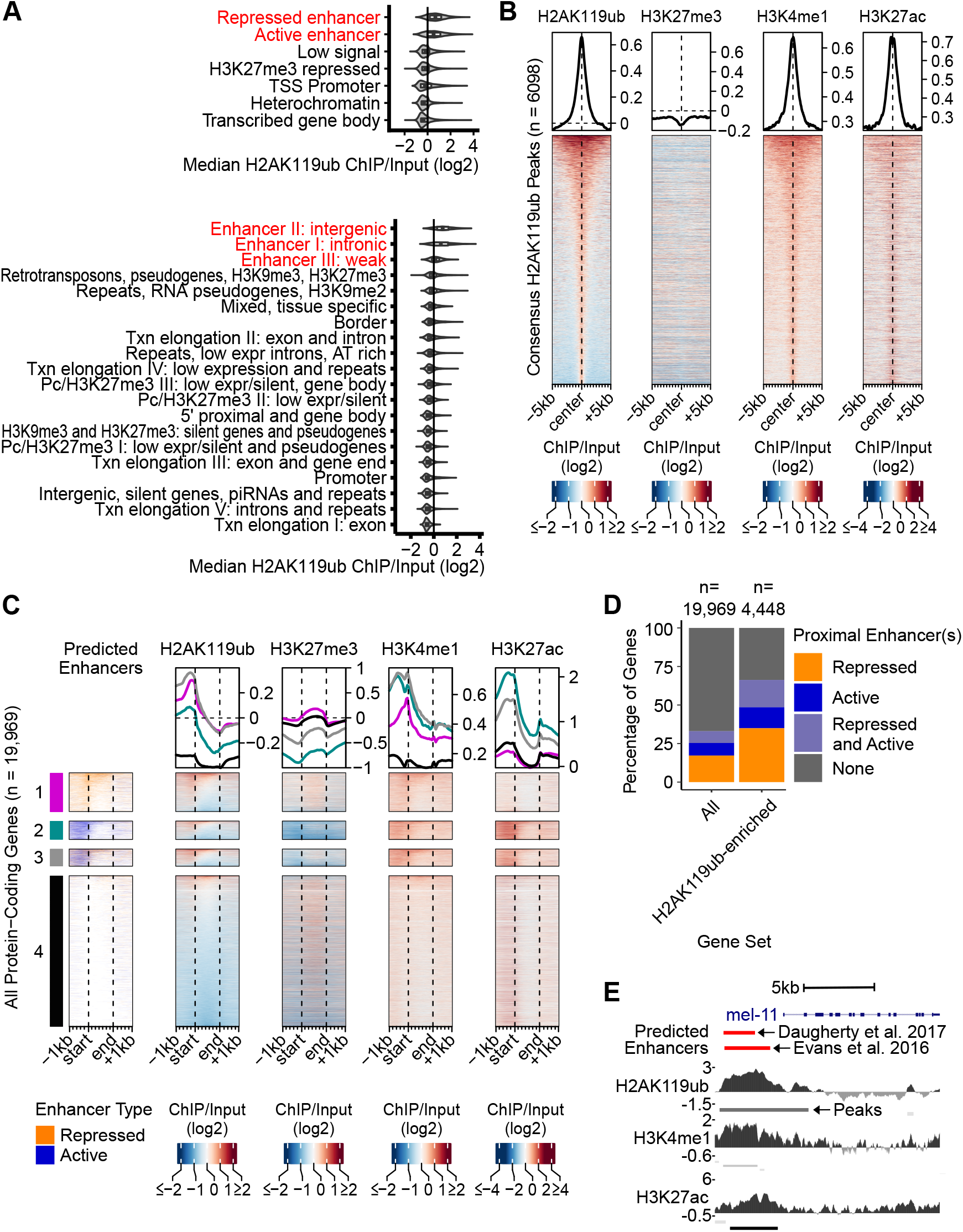
H2AK119ub is enriched at enhancer-like domains. **A.** Median H2AK119ub signal (log2-transformed ChIP/Input ratios) over chromatin state maps. The top panel includes 7 HMM-predicted chromatin states based on 8 histone modification ChIP-seq datasets (Daugherty et al. 2017). The bottom panel includes 20 HMM-predicted chromatin states based on 17 histone and histone modification ChIP-chip/ChIP-seq datasets (Evans et al. 2016). Chromatin states are arranged in order of decreasing H2AK119ub signal. **B.** Heatmaps of H2AK119ub, H3K27me3, H3K4me1, and H3K27ac (log2-transformed ChIP/Input ratios) at H2AK119ub consensus peaks. Consensus peaks are those called in two H2AK119ub ChIP-seq biological replicates. **C.** Heatmaps of predicted enhancer positions (Daugherty et al. 2017), H2AK119ub, H3K27me3, H34K4me1, and H3K27ac (log2 transformed ChIP/Input ratios) at all *C. elegans* protein-coding genes and their flanking regions. Genes are grouped into 4 clusters, depending on their annotation with a predicted enhancer region(s): cluster 1: repressed enhancers, cluster 2: active enhancers, cluster 3: both repressed and active enhancers, cluster 4: no enhancer. **D.** Quantification of the proportion of all protein-coding genes or protein-coding genes associated with an H2AK119ub consensus peak associated with the indicated type of predicted enhancer. **E.** UCSC Genome Browser view of a representative genomic region with predicted enhancer regions (Daugherty et al. 2017; Evans et al. 2016), log2 transformed H2AK119ub, H3K4me1, and H3K27ac ChIP/Input ratios and MACS2-predicted peaks. Darker colours for the predicted peaks indicate higher confidence.

To relate the enrichment of H2AK119ub at predicted enhancers to gene regulation, we assigned the predicted enhancers to the nearest gene, with a preference for a downstream gene, and compared the proportion of all protein-coding genes with a proximal enhancer to the proportion of genes with a proximal enhancer and an H2AK119ub peak. Approximately one-third of all protein-coding genes were assigned nearby predicted enhancers (33%, Daugherty et al. 2017; 27%; Evans et al. 2016) (Fig. 6C,D, Supplemental Fig. S9). In contrast, approximately two-thirds of genes with an H2AK119ub peak were assigned a proximal enhancer (66%, Daugherty et al. 2017; 59%; Evans et al. 2016) (Fig. 6C,D, Supplemental Fig. S9), a significantly higher proportion than all genes (p < 2.2e-16, Fisher’s exact test). In addition, enhancer-associated H3K4me1 and H3K27ac are frequently co-enriched with H2AK119ub at these genes (Fig. 6C, Supplemental Fig. S9A). For example, upstream of a representative gene, *mel-11,* are co-occurring peaks of H2AK119ub, H3K4me1, and H3K27ac that overlap with predicted enhancers (Fig. 6E). Together, these findings support a role for H2AK119ub, potentially in cooperation with H3K4me1 and H3K27ac, at enhancer-like domains in *C. elegans*.

To determine if the activity of H2AK119ub-enriched enhancers are impacted by the loss of *mig-32(n4275)* or *spat-3(mgw26)*, we determined if the differentially expressed genes are associated with H2AK119ub-enriched active or repressed enhancers (Fig. 7). The active or repressed enhancer states corresponds to enhancers with H3K27ac or weak H3K27me enrichment, respectively (Daugherty et al. 2017). Sixty percent of H2AK119ub-enriched differentially expressed genes have a nearby predicted enhancer, which is significantly higher than the proportion of all differentially expressed genes assigned enhancers (35%, Daugherty et al. 2017; 30%; Evans et al. 2016, p < 2.2e-16, Fisher’s exact test) (Fig. 7A,B, Supplemental Fig. S10A,B). Furthermore, H2AK119ub-enriched differentially expressed genes are 1.94-fold more likely to have an associated repressed enhancer compared to all differentially expressed genes (p < 2.2e-16, Fisher’s exact test) (Fig. 7B). Repressed enhancers associated with differentially expressed genes are more highly enriched for H2AK119ub compared to those associated with non-differentially expressed genes (p < 4.1e-05, Mann-Whitney U Test), and 72% (108/150) of the H2AK119ub-enriched differentially expressed genes with a repressed enhancer are up-regulated in the H2AK119ub-deficient mutants (Fig, 7A,C). In contrast, the active enhancers (Daugherty et al. 2017) that are associated with differentially expressed genes have on average less H2AK119ub enrichment than those associated with non-differentially expressed genes (p < 8.9e-03, Mann-Whitney U Test), and 77% (74/95) of the differentially expressed genes with a nearby active enhancer are down-regulated in the *mig-32* and *spat-3* mutants (Fig. 7A,C). Together, this suggests that the PRC1-like complex regulates enhancer activity, with a more prominent role for H2AK119ub at repressed versus active enhancers.

**Figure 7.**
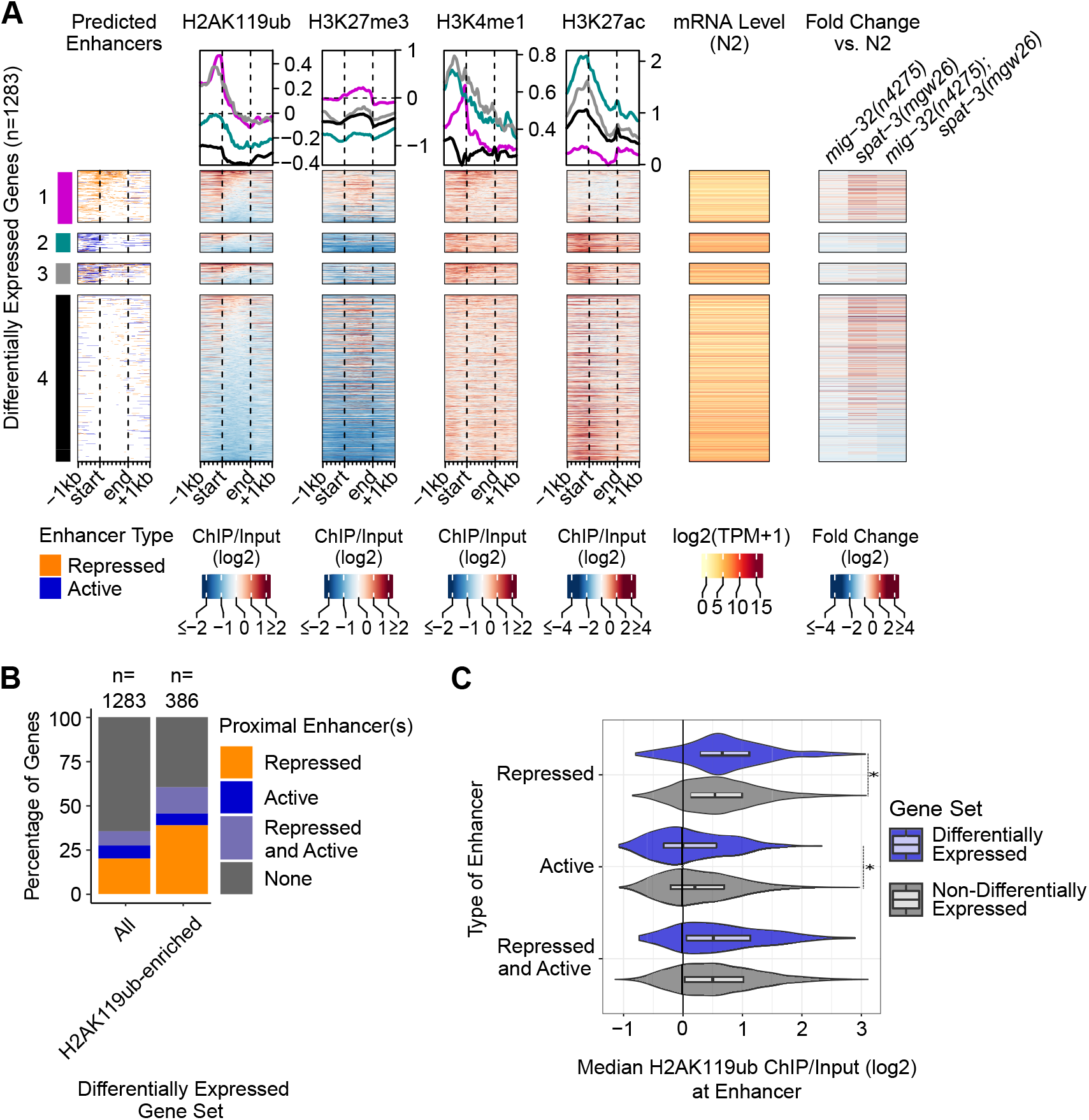
Differential expression of genes with proximal H2AK119ub-enriched predicted enhancers. **A.** Heatmaps of predicted enhancer positions (Daugherty et al. 2017), H2AK119ub, H3K27me3, H3K4me1, and H3K27ac (log2 transformed ChIP/Input ratios) at the differentially expressed protein-coding genes and their flanking regions. Genes are grouped into 4 clusters, depending on their annotation with a predicted enhancer region(s): cluster 1: predicted repressed enhancers, cluster 2: predicted active enhancers, cluster 3: both predicted repressed and active enhancers, cluster 4: no enhancer. **B.** Quantification of the proportion of all protein-coding genes or protein-coding genes associated with an H2AK119ub consensus peak associated with the indicated type of predicted enhancer. **C.** Median H2AK119ub signal (log2-transformed ChIP/Input ratios) at predicted enhancers associated with differentially expressed genes and non-differentially expressed genes. Asterisks indicate a significant difference in median H2AK119ub levels (p < 0.01, Mann-Whitney U test).

## Discussion

In this study, we have shown that H2AK119ub and H3K27me3, two histone modifications that are historically thought to cooperatively mediate gene repression, are not generally enriched at the same genomic locations and are associated with mostly distinct sets of protein-coding genes in *C. elegans* embryos. The levels of H2AK119ub and H3K27me3 are not dependent on one another being present, suggesting that they are regulated independently. We have established that, surprisingly, H2AK119ub is enriched at enhancer-like domains, with strong co-enrichment of H3K4me1 and H3K27ac. Many differentially expressed genes in *mig-32* and *spat-3* mutants have H2AK119ub-enriched promoters and enhancer elements. Among these H2AK119ub-enriched differentially expressed genes, some are normally H3K27me3-repressed whereas others are highly expressed. Given these results, we propose that H2AK119ub plays a dual role in the regulation of gene expression, involved in regulation of both H3K27me3-repressed and enhancer chromatin states.

### Interdependent regulation of H2AK119ub and H3K27me3

In some developmental contexts, the levels of H2AK119ub and H3K27me3 are highly interdependent, however, whether this is a widely conserved feature across species is uncertain. In the interdependent recruitment model, PRC1 complexes are recruited to sites of H3K27me3 and PRC2 complexes are recruited to sites of H2AK119ub by subunits that can recognize and bind each mark (Cooper et al. 2016; Chittock et al. 2017; Kuroda et al. 2020; Kasinath et al. 2021). Therefore, loss of either histone modification can impede recruitment and accumulation of both complexes, as well as their corresponding histone modifications, at target regions (Tamburri et al. 2020; Hickey et al. 2022). Indeed, disruption of PRC1 results in a global reduction of H3K27me3 enrichment and PRC2 component binding in mouse embryonic stem cells, early zebrafish embryos and *Drosophila* embryos (Fursova et al. 2019; Tamburri et al. 2020; Dobrinić et al. 2021; Hickey et al. 2022; Pengelly et al. 2015). Similarly, loss of vPRC1 or PRC2 components leads to depletion of both H2AK119ub and H3K27me3 at a subset of Polycomb targets, including the inactive X chromosome, in mouse embryoid bodies and embryonic fibroblasts (Almeida et al. 2017; Colognori et al. 2019; Sugishita et al. 2021). In addition, maternal depletion of either Polycomb complex disrupts both histone modifications at maternal imprinting sites in early mouse embryos (Chen et al. 2021; Mei et al. 2021).

In contrast to the highly interdependent levels of H2AK119ub and H3K27me3 described above, we observed that, in *C. elegans*, the bulk levels of these two histone modifications are regulated independently and the distribution of H3K27me3 was not disrupted in H2AK119ub-deficient animals, with the caveat that our study only investigates one time point in development. No homologs for an H2AK119ub-binding PRC2 component have been identified in *C. elegans*, nor any H3K27me3-binding PRC1 subunit homologs (Gahan et al. 2020). H3K27me3-binding proteins have been identified (Saltzman et al. 2018), however, it is not known if any function within a PRC1-like complex. Thus, one possibility is that *C. elegans* has evolved a distinct mechanism to regulate the levels and distribution of H2AK119ub and H3K27me3 that does not depend on the presence of the other complex or detection of the other histone modification. Identification of the PRC1-like and PRC2 interacting proteins will be required to elucidate this unique targeting mechanism.

We found that H2AK119ub and H3K27me3 are associated with mostly distinct gene sets and that H3K27me3 but not H2AK119ub is enriched at silenced genes. Nonetheless, we also identified a co-enriched subset of genes. Among these H2AK119ub and H3K27me3 co-enriched genes, the majority were not differentially expressed in H2AK119ub-deficient mutant embryos. These genes are potential candidates for redundant regulation by H2AK119ub and H3K27me3. Indeed, PRC1 and PRC2 complexes can act in parallel to repress lineage-specific genes and transcription factors in mammals (Leeb et al. 2010; Zepeda-Martinez et al. 2020; Cohen et al. 2021). Combinatorial depletion of *C. elegans* PRC1-like and PRC2 complexes will be required to determine the target genes and roles of such functional redundant gene regulation.

### Control of neuronal gene expression by Polycomb Repressive Complexes

PRC1 activity within neuronal tissues is crucial for proper regulation of cell fate and gene expression patterns, but its impact and cooperation with PRC2 can be developmental stage- and cell type-specific. For example, in mammalian neuronal stem and progenitor cells, PRC1 functions together with PRC2 to maintain neuronal differentiation factors in a poised state and is critical for maintenance of proliferative and differentiation capacity (Molofsky et al. 2003; Zencak et al. 2005; Fasano et al. 2007; Román-Trufero et al. 2009; Tsuboi et al. 2018; Yao et al. 2018). In contrast, in differentiating mammalian spinal motor neurons, PRC1 acts independently of PRC2 to maintain appropriate transcriptional programs of distinct neuronal subtypes (Sawai et al. 2022). In *C. elegans* neuronal progenitor cells, it was recently shown that the PRC1-like components *mig-32* and *spat-3* are involved in initiation of appropriate expression levels of lineage-specific transcription factors, and in differentiated neurons, PRC1-like components function independently of PRC2 to maintain cellular identity (Bordet et al. 2022). In addition, Polycomb complexes are required in various tissues to repress non-cell lineage genes, and disruption of the complex activity can promote inappropriate cell fates and gene expression patterns (Chiacchiera et al. 2016; Cohen et al. 2018).

We identified H2AK119ub-enriched and H2AK119ub and H3K27me3 co-enriched genes involved in neurogenesis, neuron differentiation, and trans-synaptic signaling that are mis-regulated in the *mig-32* and *spat-3* mutants. Misregulation of these neuronal genes may contribute to the observed abnormal neurodevelopment defects in PRC1-like mutants (Karakuzu et al. 2009; Pierce et al. 2018), however, whether this is due to gene misregulation within neuronal or non-neuronal tissues is unknown. One possibility is that the intrinsic and extrinsic factors critical for neuronal development, or the transcription factors that regulate them, are mis-regulated in *mig-32* and *spat-3* mutants, leading to inappropriate guidance cues. Since regulators of neuronal development can originate in both neuronal and non-neuronal tissues (Godini et al. 2022), appropriate PRC1-like activity may be needed in either or both tissue types. Embryos include both neural progenitor cells and differentiated neurons (Sulston 1983), and therefore, our embryonic RNA-seq data cannot differentiate between the activity of the PRC1-like complex in these different cell types. Based on the known roles of PRC1 in mammals, *Drosophila* and zebrafish, the *C. elegans* PRC1-like complex may be functioning with PRC2 to establish appropriate expression patterns in undifferentiated cells and functioning independently of PRC2 to maintain cell fates in differentiated neurons. Therefore, investigation of developmental stage-specific and cell type-specific activities will be integral to understanding the mechanism by which the PRC1-like complex guides *C. elegans* development.

### A role for H2AK119ub on the X-chromosome?

Chromatin-based mechanisms play conserved roles in regulating sex chromosomes across species. Here we find an enrichment of H2AK119ub coverage across the X-chromosome and H2AK119ub peaks at X-linked genes, thus implicating the *C. elegans* PRC1-like complex in dosage compensation. In *C. elegans* hermaphrodites, the two X chromosomes are repressed by PRC2 and H3K27me3 in the germline and by the dosage compensation complex and H4K20me1 in the soma (Strome et al. 2014). Since H2AK119ub and H3K27me3 are co-enriched on the X, this chromosome may contain genomic locations that are regulated by the two marks cooperatively in the germline. The utilization of both histone modifications for repression on a chromosome-wide scale would be consistent with the importance of both PRC1 and PRC2 in mammalian X-inactivation (Grossniklaus and Paro 2014), although the specific mechanisms of X-chromosome silencing are distinct between mammals and *C. elegans*. Alternatively, H2AK119ub may be enriched on the X-chromosome in somatic cells by the dosage compensation complex (DCC). H2AK119ub could potentially function in a similar manner to, or along with, H4K20me1, which is a downstream effector of DCC binding that leads to X-linked gene silencing via chromatin compaction (Vielle et al. 2012; Kramer et al. 2015; Brejc et al. 2017). An attenuation of topologically associated domains on the X-chromosome, as is seen in H4K20me1-deficient mutants (Bian et al. 2017), in the *mig-32* or *spat-3* mutants would support a role for PRC1 and H2AK119ub in X-chromosome conformation.

### A conserved role for H2AK119ub at enhancers

We have shown that in *C. elegans* embryos, H2AK119ub is enriched at predicted enhancer regions. The differentially expressed genes in the H2AK119ub-deficient mutants that have proximal repressed enhancers tend to be up-regulated, suggesting that this histone modification may be required to limit enhancer activity. In mammalian embryonic stem cells, poised enhancers, those enriched with both H3K4me1 and repressive Polycomb-deposited H3K27me3, are important regulators of developmental genes (Rada-Iglesias et al. 2011; Zentner et al. 2011; Cruz-Molina et al. 2017; Crispatzu et al. 2021). In *C. elegans*, H2AK119ub could be contributing to a poised or bivalent enhancer state. In this model, loss of H2AK119ub would allow the originally repressed enhancers to acquire an active chromatin state, thereby leading to a corresponding increase in enhancer-driven gene expression.

In contrast, most of the differentially expressed genes with proximal active enhancers were down-regulated in the *mig-32* and *spat-3* mutants. The associated active enhancers were not strongly enriched for H2AK119ub, however, the change in expression of the proximal genes supports a change in enhancer-mediated gene regulation, potentially the result of a loss of the PRC1-like complex components themselves rather than a loss of H2AK119ub. In *Drosophila* and mouse neural progenitor cells, cPRC1 components associate with enhancers and promoters to mediate cPRC1-anchored looping, which correlates with active transcription (Loubiere et al. 2020). A similar pattern has been observed in breast cancer and leukemia cell lines in which RING1B in the context of a cPRC1 complex displays binding to enhancers, resulting in an increase in transcriptional activity and chromatin accessibility (Chan et al. 2018; Zhang et al. 2020, 2021). If a similar mechanism of enhancer regulation is mediated by the *C. elegans* PRC1-like complex, the loss of either MIG-32 or SPAT-3 could impede enhancer-driven gene expression.

### Potential canonical and variant PRC1 functions in *C. elegans*

Multiple PCGF proteins are encoded in the mouse, human, *Drosophila*, and zebrafish genomes. Each PCGF defines a distinct PRC1 complex with a unique combination of accessory subunits and associated activities (Scelfo et al. 2019; Gahan et al. 2020). Currently, MIG-32 is the only putative functional PCGF homolog identified in *C. elegans* (Karakuzu et al. 2009; Gahan et al. 2020). However, it is not known if it is the only PCGF protein encoded in the genome or if it always functions in the same protein complex. The presence of multiple PRC1-like complexes could explain the observed H3K27me3-dependent and H3K27me3-independent activities and why fewer differentially expressed genes were observed in the *mig-32(n4275)* mutants than in the *spat-3(mgw26)* single or double mutants. SPAT-3 is a putative functional homolog of RING1A/B (Karakuzu et al. 2009; Pierce et al. 2018), an invariable PRC1 component, and therefore would be more likely to be included in multiple distinct *C. elegans* PRC1-like complexes, and consequently may have more gene regulatory targets. The majority of H2AK119ub is lost in the *mig-32(n4275)* mutants, which suggests that MIG-32 is required for H2AK119ub deposition, an activity that is attributed to vPRC1-like complexes (Chittock et al. 2017; Kuroda et al. 2020). However, MIG-32 has been shown to mediate compaction of recombinant nucleosome arrays *in vitro*, an activity that is more consistent with a cPRC1-like mechanism of gene repression (Grau et al. 2011). If MIG-32 is functioning in a cPRC1-like complex to mediate H2AK119ub-independent gene repression, it could be acting redundantly with PRC2 and H3K27me3 (Leeb et al. 2010; Cohen et al. 2021).

Using *C. elegans* embryos as a model, we have shown that H2AK119ub and H3K27me3 may regulate gene expression largely independently but could function cooperatively or redundantly at a subset of protein-coding genes. Furthermore, we have begun elucidating a previously unappreciated role for H2AK119ub at enhancers, establishing a dual role for the histone modification at two different chromatin states and expanding the repertoire by which Polycomb complexes can mediate eukaryotic genome regulation.

## Methods

### Worm maintenance

Strains were maintained on nematode growth medium (NGM) agar with *E. coli* OP50-1 as a food source as described (Stiernagle 2006) at 21°C unless specified otherwise. The N2 (Bristol) strain was used as wild-type (Brenner 1974). Strains and alleles used are listed in Supplemental Table S1.

### Synchronized worm growth

Embryos were isolated from gravid adults using standard alkaline hypochlorite treatment (“bleaching”) as described (Stiernagle 2006). After three washes in M9 buffer supplemented with 0.01% (v/v) Triton X-100 (M9/Tx), embryos were allowed to develop to L1 diapause in M9/Tx buffer overnight with aeration.

### Chromatin immunoprecipitation (ChIP)

Animals arrested at L1 diapause were prepared by bleaching as described above. Arrested L1s were counted and seeded into 40 mL of S-Complete medium (Stiernagle 2006) with OP50-1 as a food source, at a final density of ∼0.5-1.5 animals per uL. Animals were grown with aeration for 72 hours to gravid adulthood, and embryos were isolated again by bleaching. After washes with M9/Tx, embryos were processed for ChIP as described below either immediately or after a further 3 hours of growth with aeration.

Chromatin immunoprecipitation was performed essentially as described (Ercan et al. 2007; Rechtsteiner et al. 2010; Askjaer et al. 2014) with minor modifications as follows. Embryos were collected by centrifugation (1000*g*, 3 min.) and washed twice in phosphate-buffered saline (PBS) containing 0.01% (v/v) Triton X-100 (PBS/Tx). To crosslink, PBS/Tx was added to at least 80 times the pellet volume, and 37% formaldehyde (Sigma F8775) was added to a final concentration of 1.8%. Samples were incubated at room temperature with rotation for 25 minutes. Glycine was added to a final concentration of 125 mM followed by incubation for an additional 5 minutes. All subsequent steps were performed on ice using pre-chilled buffers and tubes. Embryos were washed twice with PBS/Tx as above and an equal volume of resuspension buffer (50 mM HEPES-KOH pH7.5, 150 mM NaCl, 1 mM EDTA, 0.01% Triton X-100, protease inhibitors (1 tablet per 5 mL, Roche 4693159001)) was added to the embryo pellet. The resuspended embryos were aliquoted, snap-frozen in liquid nitrogen and stored at -80°C.

Aliquots of cross-linked embryos were thawed in an ice-water bath, the volume was adjusted to 3-4 pellet volumes (PV) using resuspension buffer, and 100 uL was aliquoted to each polystyrene sonication tube (Evergreen 214-3721-010). An equal volume of resuspension buffer containing 2X detergents was added (50 mM HEPES-KOH pH7.5, 150 mM NaCl, 1 mM EDTA, 0.2% sodium deoxycholate, 0.7% sodium sarkosyl). Samples were sonicated in a Q800R3 water bath sonicator (QSonica Q800R3-110; 50% power, 20s on/20s off) for 4 minutes. Samples were gently mixed by pipetting and sonicated for an additional 4 minutes.

The lysate was transferred to an Eppendorf tube and an equal volume of resuspension buffer without detergents was added (50 mM HEPES-KOH pH7.5, 150 mM NaCl, 1 mM EDTA). The lysate was centrifuged (13,000*g*, 15 min. 4°C) and the supernatant was transferred to a new tube. Protein concentration was determined using the Bradford assay (Sigma B6916).

The supernatant was aliquoted to tubes for immunoprecipitation (IP), saving 20% of the volume of one IP tube as the ‘input’ control, which was stored at -20°C. Triton X-100 (10%, v/v) was added to a final concentration of 1% (v/v). This step was omitted for anti-H3K27me3 (CST 9733, C36B11), for which this detergent led to increased background (Saltzman, unpublished observations). See Supplemental Table S2 for antibody details. After antibody addition, samples were incubated overnight at 4°C with rotation. Protein G dynabeads (Invitrogen 10003D; 7.5 uL per IP) were pre-washed twice with resuspension buffer and added to IPs, which were then incubated as above for 2 hours. Beads were collected on a magnetic stand (Permagen MSR812) and washed as follows, with each wash incubated for 5 minutes at 4°C with rotation: 2X with FA-150 (50 mM HEPES-KOH pH 7.5, 150 mM NaCl, 1 mM EDTA, 1% Triton X-100, 0.1% Sodium deoxycholate), 1X with FA-1M (FA-150 with 1 M NaCl), 1X with FA-0.5M (FA-150 with 0.5 M NaCl), 1X with TE-LiCl in a new tube (10 mM Tris-Cl pH 8, 1 mM EDTA, 250 mM LiCl, 1% IGEPAL CA-630, 1% sodium deoxycholate), 2X with TE+ (10 mM Tris-Cl pH 8, 1 mM EDTA, 50 mM NaCl, 0.005% IGEPAL CA-630).

Elution was performed in a new tube at 65°C for 15 minutes using 50 uL of ChIP elution buffer (10 mM Tris-Cl pH 8, 1 mM EDTA, 200 mM NaCl, 1% SDS). Elution was repeated and the supernatants were pooled. The ‘Input’ aliquot was thawed and brought to 100 uL with ChIP elution buffer. Input and IP samples were treated with RNase A (16.5 ug, Sigma R4642) for 1 hour at 37°C and proteinase K (40 ug, Bioline BIO-37084) for 2 hours at 55°C. Crosslinks were reversed by incubation overnight at 65°C. DNA was isolated using a QIAquick PCR purification kit (Qiagen 28104). Input DNA was quantified using NanoDrop spectrophotometer or Qubit Fluorometer (ThermoFisher).

### ChIP-seq library preparation and ChIP-qPCR

Libraries were prepared with the NEBNext Ultra II DNA Library Prep Kit for Illumina (NEB E7645S) according to the manufacturer’s instructions, using 25-50 ng of input DNA or half of ChIP DNA. Library concentration was quantified using NEBNext Library Quant Kit for Illumina (NEB E7630). Pooled libraries were sequenced using NextSeq500 in 75 bp paired-end mode. Details of read counts are in Supplemental Table S2.

For qPCR, input DNA samples were diluted 1:50 and ChIP samples were diluted 1:4. qPCR was performed with iTaq Universal SYBR Green Supermix (Biorad 1725121) in a 10 uL reaction. Primer efficiency was assessed by a 4-fold dilution series of input DNA.

### Analysis of C. elegans ChIP-seq data

The raw sequencing reads were trimmed for low-quality base calls, with a phred33 < 22, Illumina adapter sequences using a stringency of 12, and poly-G sequences (defined as 13 Gs in a row) using a stringency of 8 with TrimGalore v0.6.5 (Andrews 2010; Martin 2011; Krueger 2015). The trimmed reads were then aligned to the *Caenorhabditis elegans* ce11 genome using Bowtie2 v2.3.4.3 (Langmead and Salzberg 2012) with the maximum fragment size parameter set to 2000. Alignments were filtered by removing duplicates and multi-mapping reads with a MAPQ < 42, using Samtools v1.11 (Li et al. 2009). Peaks were called with MACS2 v2.2.7.1 (Zhang et al. 2008) with -*g* option set to ce, -*f* option set to either BAMPE for paired-end libraries or BAM for single-end libraries, and the --*broad* option for all modifications except for H3K4me1, H3K4me3, and H3K27ac. Coverage files were prepared with the BedTools v2.29.0 (Quinlan and Hall 2010) genomecov function with the -*bga* option. For paired end data the *-pc* option was used. For single end data the -*fs* option was set as the fragment size predicted using SPP v1.15.4 (Kharchenko et al. 2008). Log2 transformed ChIP/Input ratios were calculated per base with deepTools v3.5.0 bigwigCompare (Ramírez et al. 2014) with a pseudocount of 1.

Peaks that overlap with blacklisted regions, where there is artificially high signal, were removed (Amemiya et al. 2019). Consensus peaks were defined as peaks from two ChIP-seq replicates that overlap. ChIP-seq peaks were called as overlapping if at least 50% of the peak region overlapped in two samples. The overlap was determined using the BedTools v2.29.0 (Quinlan and Hall 2010) intersect function with the -*f* option set to 0.5 and the -*u* option. This function was then repeated with the -*a* and -*b* input files switched. To identify common peaks, overlapping peak sets were combined with the BedTools merge function with -*c* option set to 4,5,6 (name, score, and strand BED fields) and -*o* option set to collapse, mean, distinct. Unique (non-overlapping) peaks for input file A were determined with the BedTools intersect function with the -*f* option set to 0.5 and the *-v* option included. Unique peaks for input file B were determined with the same function but -*a* and -*b* input files switched.

### Assigning ChIP-seq peaks and predicted enhancer regions to protein-coding genes

ChIP-seq peaks or predicted enhancer regions (Evans et al. 2016; Daugherty et al. 2017) were assigned to the nearest gene in the Wormbase WS263 release with ChIPseeker v1.28.3 (Yu et al. 2015). The promoter region was defined as -1kb for assignment of ChIP-seq peaks. The promoter region was defined as -500bp for assignment of predicted enhancers. Gene assignments were filtered for protein-coding genes (n=20,094), and genes with no H2AK119ub or H3K27me3 ChIP or Input signal across the entire gene body, 1kb upstream, or 1kb downstream were removed (n=19,969). See Supplemental Table S3 for details of gene assignments.

### Genomic data visualization

Matrices of ChIP-seq signals were produced with either the computeMatrix function of deepTools v3.5.0 (Quinlan and Hall 2010) or the normalizeToMatrix function of EnrichedHeatmap v1.22.0 (Gu et al. 2018). The matrices used to plot the violin plots of H2AK119ub and H3K27me3 at gene promoters, were prepared with computeMatrix with the –*upstream* option set to 500 and the *–referencePoint* set to the gene start site. The matrices used to plot the violin plots of H2AK119ub at different chromatin states, were prepared with computeMatrix with the -*m* option set to 1000. computeMatrix was used with the –*binsize* set to 50000 for the chromosomal distribution heatmaps. normalizeToMatrix was used for all other matrices of ChIP-seq signal with the window size set to 50 bp, background set to 0, and the mean mode set to w0. The positions of the predicted enhancer regions (Evans et al. 2016; Daugherty et al. 2017) were visualized with matrices computed with normalizeToMatrix with the window size set to 50 bp, background set to 0, and the mean mode set to absolute. All heatmaps were made with the EnrichedHeatmap v1.22.0 (Gu et al. 2018) package, except for the heatmap of the Spearman Correlation coefficients of histone modifications across all chromosomes. For this heatmap, correlation was computed using the base R function *cor* with method set to ‘spearman’ and use set to ‘complete.obs’ and the heatmap was plotted using Pheatmap v1.0.12 (Kolde 2019). Coverage tracks and peaks were visualized with the UCSC Genome Browser (Kent et al. 2002).

### Mammalian ChIP-seq data

The fastq files for all mammalian ChIP-seq data were downloaded from Gene Expression Omnibus (GEO) (Edgar et al. 2002). Details of GEO accession and SRA accession numbers can be found in Supplemental Table S2. The raw sequencing reads were trimmed for low-quality base calls, with a phred33 < 20 and Illumina adapter sequences using a stringency of 3 with TrimGalore v0.6.5 (Andrews 2010; Martin 2011; Krueger 2015). The reads from the iPSCs and T47D cells also included the option -- *three-Prime-clip-R1 3* in the TrimGalore call. The trimmed reads were then aligned to the *Mus musculus* mm10 or the *Homo sapiens* hg38 genome using Bowtie2 v2.3.4.3 (Langmead and Salzberg 2012). All other processing is as described in ‘Analysis of *C. elegans* ChIP-seq data’ and ‘Genomic data visualization’ above.

### Protein sequence alignment

Protein sequences were obtained from the National Center for Biotechnology Information (Sayers et al. 2022) and were aligned with Clustal Omega using default settings (Sievers et al. 2011). The alignments were visualized using the ESpript server (Robert and Gouet 2014).

### Total RNA isolation

Worms were synchronized by bleaching as described above and grown to gravid adulthood on NGM agar plates with OP50-1. To collect late embryos for RNA-seq, the synchronized gravid adults were bleached 60 hours (N2) or 66 hours (*mig-32* and *spat-3* mutants) after release from L1 diapause. Embryos of all genotypes were washed three times in M9/Tx buffer and allowed to develop for 4.5 hours post bleaching in M9/Tx buffer with aeration. The developmental stages of a subset of each sample were scored with DIC microscopy before collection (Supplemental Fig. S7A). TRI Reagent (Sigma Aldrich) was added to the embryo pellet followed by 1-3 cycles of snap-freezing, thawing, and vortexing (1 minute). Following chloroform extraction, the aqueous phase was transferred to an RNA Clean and Concentrator-5 column and purified according to the manufacturer’s instructions including on-column DNaseI digestion (Zymo Research).

### RNA-seq library preparation

PolyA+ RNA was isolated from total RNA using the NEBNext Poly(A) mRNA Magnetic Isolation Module (NEB E7490) according to the manufacturer’s instructions. Libraries were prepared with the NEBNext Ultra II Directional RNA Library Prep with Sample Purification Beads (NEB E7765S) according to the manufacturer’s directions with NEBNext Multiplex Oligos for Illumina (NEB E6440). Library concentration was quantified using NEBNext Library Quant Kit for Illumina (NEB E7630). Pooled libraries were sequenced using NextSeq500 in 75 bp paired-end mode. Details of read counts are in Supplemental Table S2.

### Analysis of RNA-seq data

The raw sequencing reads were trimmed as described in ‘Analysis of *C. elegans* ChIP-seq data’ above. The trimmed reads were then aligned to the *Caenorhabditis elegans* ce11 genome using STAR v2.7.9a (Dobin et al. 2013) with default parameters and filtered for those with MAPQ 255 with Samtools v1.12 (Li et al. 2009). Mapped reads were quantified with Subread v2.0.1’s featureCounts (Liao et al. 2014). Differentially expressed genes were identified with DESeq2 v1.32.0 (Love et al. 2014) using a cutoff of a p-value ≤ 0.05, calculated with the Wald test and corrected for multiple testing using the Benjamini-Hochberg method (Benjamini and Hochberg 1995), and a fold-change of ≥ 1.5 in either direction. Heatmaps of the normalized counts and log2 fold changes of the differentially expressed genes were prepared with Pheatmap v1.0.12 (Kolde 2019). See Supplemental Table S3 for details of wild-type transcript per million counts, log2 fold changes and p-adjust values for each protein-coding gene in *mig-32* and *spat-3* mutants.

### Gene Ontology (GO) Analysis

For gene ontology analysis, the enrichGO function of clusterProfiler v4.0.5 (Yu et al. 2012) was used to search for enriched biological process terms. For protein-coding genes assigned ChIP-seq peaks, the background gene set was defined as all protein-coding genes (n=20,094). For differentially expressed genes, the background gene set was defined as all protein-coding genes that were detectable in embryos, those with a raw fragment count of at least 10 in at least one sample (n=14,652). The BioConductor org.Ce.eg.db v3.13.0 database was used and a q-value cutoff of 0.05. The resulting lists were filtered for redundant terms with clusterProfiler’s simplify function, using a similarity threshold of an adjusted p-value of 0.7 and the measurement of similarity as described (Wang et al. 2007).

### RNAi

For RNAi plates, NGM agar was supplemented with 25 ug/mL carbenicillin and 5 mM IPTG. RNAi plates were seeded with *E. coli* HT115 expressing either control dsRNA (pL4440-Dest; a kind gift from Dr. J. Reece-Hoyes, Dr. A. J. M. Walhout, UMass Chan Medical School) or *mes-2* dsRNA. The *mes-2* RNAi plasmid (pT444T-mes-2) was constructed by cloning the *mes-2* cDNA into the pL4440 derivative, pT444T (Addgene plasmid # 113081; Sturm et al. 2018). RNAi bacteria were grown overnight in LB media with 1000 ug/mL ampicillin and diluted to a relative OD600 of 2.5 before seeding. Embryos of the P0 generation were grown to gravid adulthood on RNAi plates for 73 hours. F1 embryos were isolated by bleaching as described above, allowed to develop for 4 hours and harvested for western blots. The developmental stages of a subset of the F1 embryos were scored with DIC microscopy. A subset of the F1 embryos were grown to gravid adulthood on RNAi plates for 73 hours and scored for fertility defects.

### Western Blot

Late embryos were prepared as described above (see Total RNA isolation) except that the embryos were harvested 5 hours after bleaching or were harvested as described following RNAi treatment. Embryos were washed once in resuspension buffer (see ChIP), snap-frozen on dry ice in resuspension buffer, and thawed on ice. An equal volume of resuspension buffer with 2X detergents (see ChIP) was added and samples were sonicated in a Q800R3 water bath sonicator (QSonica Q800R3-110; 30% power, 15s on/45s off) for 2 minutes. The lysate was centrifuged (13,000*g*, 15 minutes at 4°C) and the supernatant was transferred to a new tube. Protein concentration was determined by Bradford assay (Sigma-Aldrich B6916). Lysates were resolved on a 4-12% Bis-Tris NuPage gel (Invitrogen), transferred to nitrocellulose, and probed with an anti-H2AK119ub antibody (1:2000, rabbit monoclonal D27C4, 8240; Cell Signaling Technology), or with an anti-H3K27me3 antibody (1:1000, rabbit monoclonal C36B11, 9733; Cell Signaling Technology). The secondary antibody (goat, HRP-conjugated anti-rabbit IgG, 7074; Cell Signaling Technology) was used at a dilution of 1:3000. Chemiluminescent signal was visualized using a ChemiDoc MP imaging system (Bio-Rad). Following stripping (15 minutes in 2% SDS, 0.1 M beta-mercaptoethanol pre-heated to 65°C), blots were re-probed with an anti-histone H3 antibody (1:6000, rabbit polyclonal ab1791; Abcam) as a loading control.

## Data access

All raw and processed sequencing data generated in this study have been submitted to the NCBI Gene Expression Omnibus (GEO; https://www-ncbi-nlm-nih-gov/geo/) under accession number GSE222475.

## Acknowledgements

We gratefully acknowledge Dr. E. Campos. Dr. J. Mitchell (University of Toronto), and members of the Saltzman lab for critical reading of the manuscript. We thank E. McCartney for cloning the *mes-2* RNAi construct. Illumina sequencing was performed by the Donnelly Sequencing Centre. This work was supported by a Natural Sciences and Engineering Research Council of Canada (NSERC) Discovery Grant (RGPIN-2019-06843) and a Canadian Institutes of Health Research Project Grant (PJT-175245) to A.L.S. K.M. was supported by an NSERC-CGS-M and an NSERC-PGS-D. D.F. and L.T. were supported by University of Toronto Excellence Awards. Some strains were provided by the CGC, which is funded by NIH Office of Research Infrastructure Programs (P40 OD010440).

